# Discovery of Inhibitors Targeting the RNA-Dependent RNA Polymerase of the Zika Virus

**DOI:** 10.1101/2025.07.15.665018

**Authors:** Vanessa Aitken, Anthony F. T. Moore, Manuel Terrazas-López, Salvador Rodarte, Eda Koculi

## Abstract

The Zika virus (ZIKV) is a Orthoflavivirus linked to several neurological and developmental diseases, and it remains a significant global health threat for which no treatments or vaccines are currently available. In this study, we optimized the malachite green colorimetric assay, previously used to measure the RNA-dependent RNA polymerase (RdRp) activity of the hepatitis C and foot-and-mouth disease viruses, and applied this assay to identify inhibitors of ZIKV RdRp, an enzyme essential for viral replication. Inhibitor screening was conducted using 204 compounds from the Natural Product Set IV of the National Cancer Institute Developmental Therapeutics Program. We identified two compounds, purpurogallin and digallic acid, as inhibitors of ZIKV RdRp. Future studies will characterize the potency and elucidate the mechanisms of action of these compounds.

## INTRODUCTION

Zika virus (ZIKV) is a positive-sense, single-stranded RNA virus belonging to the *Orthoflavivirus* genus, which can, in certain cases, cause neurological and developmental disorders ^1–3^. While the primary route of ZIKV transmission is through mosquito bites, it can also be transmitted from mother to fetus and via bodily fluids ^1–2, 4^. Recently, more than 92 countries have experienced a resurgence of ZIKV outbreaks ^4^. Moreover, rising global temperatures are expected to increase the risk of ZIKV infections, with an estimated 1.3 billion people projected to be at risk by 2050 ^5–6^. To date, treatment and vaccines for ZIKV are not available ^2^. Therefore, the development of new therapeutic agents to treat ZIKV infections is of significant public health interest ^2^. In this study, we optimized a previously used malachite green assay and used it to screen for small-molecule inhibitors of the RNA-dependent RNA polymerase (RdRp) using the Natural Product Set IV from the National Cancer Institute’s Developmental Therapeutics Program ^7–13^.

The ZIKV genome consists of single-stranded RNA that functions as mRNA, from which proteins required for viral replication and propagation are translated ^1–2^. This RNA is translated by the host cellular machinery into a single polypeptide chain comprising all ten viral proteins: C, preM, E, NS1, NS2A, NS2B, NS3, NS4A, NS4B, and NS5 ^1–2, 14^. The polypeptide is then cleaved by viral and cellular proteases to produce three structural proteins—C, preM, and E—which form the viral shell—and seven nonstructural proteins—NS1, NS2A, NS2B, NS3, NS4A, NS4B, and NS5— which are involved in viral replication and immune evasion ^14–15^. NS5 contains two domains connected by a linker: an N-terminal methyltransferase domain and a C-terminal RdRp domain ^16^. RdRp is responsible for replicating the ZIKV RNA genome and is essential for viral replication ^16^. As RdRp is encoded by the virus and not present in humans, and because it plays a critical role in the viral life cycle, it represents an attractive therapeutic target for treating ZIKV infections with potentially minimal off-target effects.

Previously, a malachite green high-throughput screening (HTS) assay was employed to measure RdRp activity from hepatitis C and foot-and-mouth disease viruses ^8^. The malachite green assay is a colorimetric assay that detects the complex formed between malachite green, molybdate, and inorganic phosphate ^7, 9–13^. However, RdRp from hepatitis C and foot-and-mouth disease viruses, and polymerases in general, produce pyrophosphate during their reaction instead of inorganic phosphate ^8^. Thus, in the previous study, the pyrophosphatase enzyme, which converts pyrophosphate to inorganic phosphate, was included in the RdRp reaction buffers ^8^.

In order to search a large library of compounds and accurately determine their modulation of RdRp activity, it is important to design a HTS assay that is reproducible from day to day and has a large dynamic range. While the malachite green assay previously employed for hepatitis C and foot-and-mouth disease virus RdRp was elegantly designed and executed, key performance metrics such as dynamic range (Z′) and signal window (SW) were not systematically evaluated ^8, 17^. In this study, we adapted the previously used malachite green assay and performed statistical analysis using Z′ and SW parameters ^17^. We then used the optimized assay to screen for natural product inhibitors of Zika virus RdRp and identified two compounds—purpurogallin and digallic acid— that modulate ZIKV RdRp activity. In its current form, the malachite green assay described here can be used to assess the activity of both DNA and RNA polymerases and to identify small-molecule modulators of this important class of enzymes in a high-throughput manner.

## MATERIALS AND METHODS

### Plasmid Construction and Protein Expression

RdRp from the ZIKV Puerto Rico strain PRVABC59 was inserted into the pET His6 Sumo TEV LIC (Novagen) by the Protein Production and Analysis Facility at the Sanford Burnham Prebys Medical Discovery Institute in Florida. The final protein construct consisted of a His-tag followed by a SUMO sequence, a TEV protease cut site, eight of the ten amino acid residues of the linker region separating the methyltransferase from RdRp, and the entire sequence of RdRp. The SUMO sequence has been shown to enhance the translation and solubility of proteins expressed in bacteria^18^. The linker region connecting the methyltransferase and RdRp was retained because this region has been shown to regulate RdRp activity ^16^. The His-tag and SUMO sequence were not cleaved. Thus, the protein assay described here was performed with the RdRp construct containing both the His-tag and SUMO sequence. The inorganic pyrophosphatase from *Saccharomyces cerevisiae* was purchased from Fisher Scientific

### RdRp Expression and Purification

*Escherichia coli* (*E. coli*) BL21 CodonPlus (DE3) cells were transformed with the pET-His₆-SUMO-TEV-LIC vector containing the RdRp coding sequence. The cells were grown in Luria Broth medium supplemented with 30 μg/mL kanamycin and 34 μg/mL chloramphenicol at 37 °C until reaching an optical density at 600 nm (OD_600_) of approximately 0.6. Protein expression was induced with 500 μM isopropyl β-D-1-thiogalactopyranoside, and the cultures were incubated for an additional 16 hours at 20°C. Cells were harvested by centrifugation.

The cell pellets were resuspended in lysis buffer consisting of phosphate-buffered saline (PBS: 137 mM NaCl, 2.7 mM KCl, 10 mM Na₂HPO₄, 1.8 mM KH₂PO₄), supplemented with 5 mM β-mercaptoethanol (BME) and 1 mM phenylmethylsulfonyl fluoride (PMFS). Cells were lysed by sonication on ice using a Branson Digital Sonifier 450 equipped with a 0.5 mm diameter tip using six 30 second pulses in ice. The lysate was cleared by centrifugation at 11,648 x g for 60 minutes at 4 °C.

The clarified supernatant was loaded onto an 8 mL Ni-NTA affinity column pre-equilibrated with PBS containing 5 mM BME, 1 mM PMFS and 20 mM imidazole. The protein was eluted using PBS supplemented with 5 mM BME, 1 mM PMFS and 500 mM imidazole. Fractions containing the RdRp protein were pooled and diluted 1:10 with a buffer consisting of 25 mM Tris-HCl (pH 8.0) and 5 mM BME.

Subsequently, the protein sample was loaded onto a 5 mL heparin affinity column (HiTrap Heparin HP, GE Healthcare Life Sciences). Heparin columns selectively bind nucleic acid–binding proteins and compete with nucleic acids for protein interaction sites ^19^. Therefore, this step facilitated both the further purification of the RdRp protein and the removal of any bound nucleic acids. The column was equilibrated and washed with a buffer consisting of 25 mM Tris-HCl (pH 8.0) and 30mM NaCl and 5 mM BME. RdRp was eluted using a linear gradient of heparin elution buffer consisting of 25 mM Tris-HCl (pH 8.0), 1000 mM NaCl, and 5 mM BME.

To further remove undesired proteins and RdRp aggregates, fractions containing RdRp were pooled, concentrated, and loaded onto a size-exclusion chromatography column (Superdex 200 10/300 GL, GE Healthcare). The column was equilibrated with gel-filtration buffer consisting of 25 mM Tris-HCl (pH 8.0), 100 mM KCl, and 2 mM dithiothreitol (DTT). Fractions containing RdRp were combined and concentrated using 10 kDa molecular weight cutoff centrifugal filters (Amicon Ultra Ultracel-10K, Millipore). Finally, purified RdRp was flash-frozen in small aliquots and stored at -80 °C until use.

### Malachite Green Assay for Screening Inhibitors of ZIKV RdRp

The RNA polymerase reaction conditions were adapted from a previous study ^7^. Pyrophosphate release during the RdRp reaction was measured using the malachite green/molybdate colorimetric assay ^7^. Because the malachite green/molybdate assay specifically detects inorganic phosphate but not pyrophosphate, inorganic pyrophosphatase was added to the reaction mixture to convert pyrophosphate to inorganic phosphate, enabling accurate quantification ^7–9^.

The RNA polymerase reaction initiation complex was formed by annealing poly(C) RNA molecules (200–250 nucleotides in length) with a 10-nucleotide-long poly(G) RNA primer ^20^. The poly(C) RNA was purchased from Sigma-Aldrich and further purified by polyacrylamide gel electrophoresis ^12, 21–23^. The HPLC-purified poly(G) oligonucleotides were obtained from Integrated DNA Technologies (IDT). Annealing was performed by mixing the two RNA strands in annealing buffer containing 10 mM Tris-HCl (pH 7.5), 100 mM NaCl, and 0.2 mM EDTA-KOH (pH 8.0). The mixture was incubated at 95 °C for 2 minutes and then allowed to slowly cool to room temperature to promote proper duplex formation ^12^.

The RdRp reaction was performed in a 384-well plate at 22°C, in 12.5 µL reaction mixture consisting of 5 mM Tris-HCl pH 7.5, 7 mM NaCl, 8 mM MnCl_2_, 10% glycerol, 0.01% Triton X-100, 10 mM DTT, 0.05 mU/µL inorganic pyrophosphatase, 0.5 mU/µL SUPERase·In, 500 nM annealed RNA, 0.5 mM GTP·Mn, 0 to 200 nM RdRp, and 1% DMSO ^20^. Reactions for compound screening also contained 10 µM of compound. The polymerase reactions were initiated by the addition of GTP·Mn, allowed to continue for 30 to 90 minutes and subsequently quenched with 10 mM EDTA, which sequesters Mn^2+^ and halts the RdRp reaction ^12, 20^.

To detect orthophosphate generated by pyrophosphatase activity, 75 µL of malachite green– molybdate reagent was added to the RdRp reaction mixture. After 1 minute, 10 µL of 34% sodium citrate was added to stop the reaction and stabilize the color complex. The mixture was then incubated at 22 °C for 50 minutes before measuring absorbance at 625 nm using a Molecular Devices SpectraMax Plus 384 spectrophotometer and microplate reader. The malachite green– molybdate reagent was prepared as previously described ^11–12^.

## RESULTS AND DISCUSSION

### Optimization of the Malachite Green Assay for ZIKV RdRp Inhibitor Screening

The malachite green assay quantifies orthophosphate by detecting the green-colored complex formed between orthophosphate, malachite green, and molybdate ^9–13^. However, during double-stranded RNA synthesis, the RdRp releases pyrophosphate as a byproduct ^8^. To enable the detection of pyrophosphate using the malachite green assay, inorganic pyrophosphatase—which converts pyrophosphate to orthophosphate—was included in the reaction mixture ^7–8^. We aimed to determine the RdRp reaction and assay conditions that yield reliable, proportional, and accurate measurements. Therefore, we evaluated the linearity of the RdRp reaction, as measured by the malachite green assay, with respect to both time and RdRp concentration. Our results demonstrate that the assay response remains linear for at least 90 minutes (Figure 1). Based on these findings, we selected a 75-minute reaction time with 200 nM RdRp to maximize signal intensity while maintaining assay linearity.

**Figure 1:**
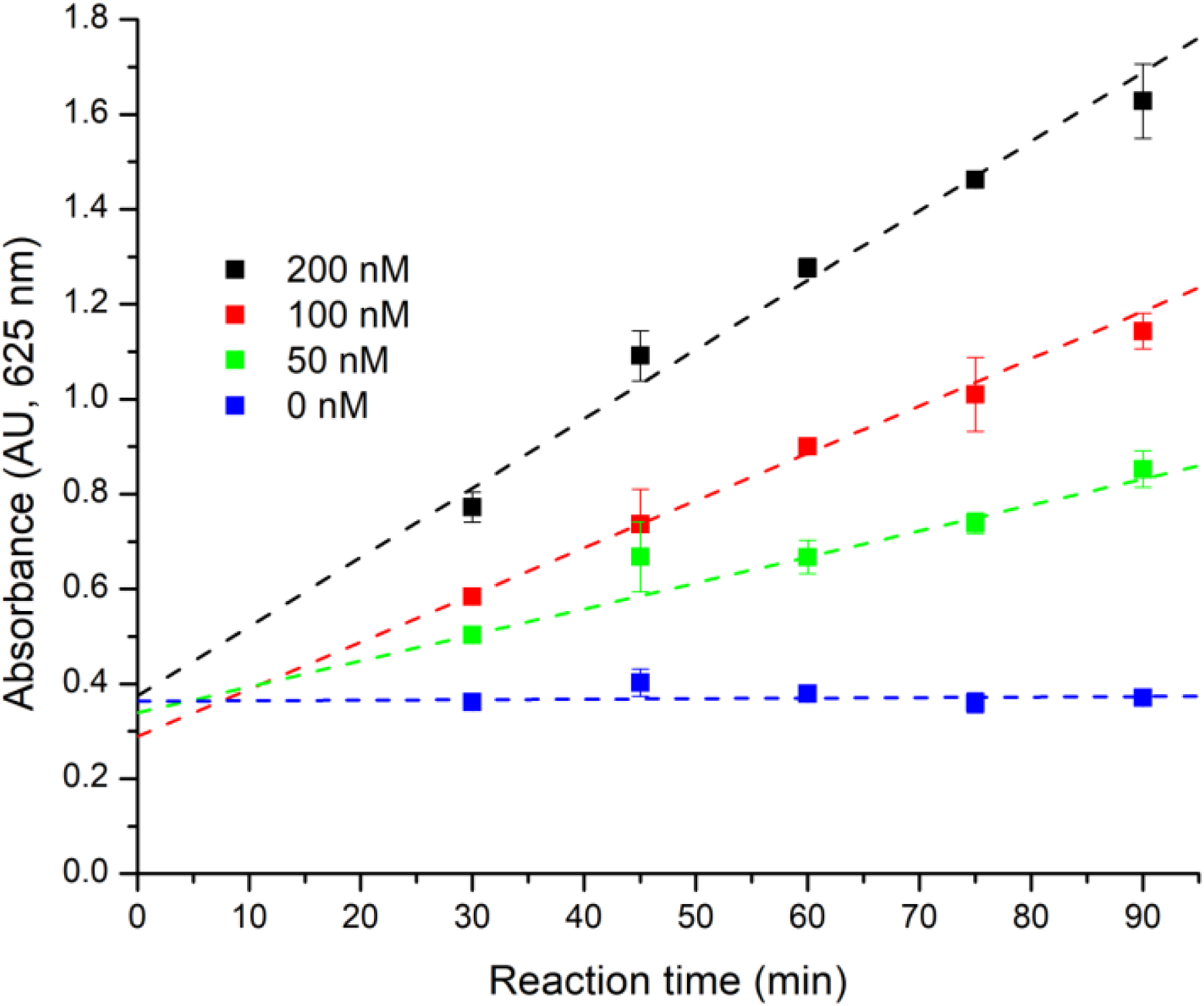
The malachite green assay used to measure RdRp activity is linear across a range of enzyme concentrations and reaction times. Dotted lines represent linear fits of the data for each RdRp concentration over increasing reaction times. Assays performed in the absence of RdRp are shown as blue squares; with 50 nM RdRp as green squares; 100 nM RdRp as red squares; and 200 nM RdRp as black squares. Data points represent the mean of three independent experiments; error bars indicate standard deviations. Based on these results, a 75-minute reaction with 200 nM RdRp was selected for inhibitor screening to maximize signal while maintaining assay linearity.

### Robustness, Reproducibility, and Dynamic Range of the Malachite Green Assay Employed in This Study

HTS robustness testing ensures that minor variations in experimental conditions across different days do not compromise the accuracy of hit detection, either by misidentifying false positives as true hits or by failing to detect true positives. These evaluations are essential for maintaining consistency and reliability in assay performance, ensuring that inter-day variability remains minimal and does not interfere with the detection of inhibitory activity. To assess the robustness of the malachite green assay used for evaluating ZIKV RdRp activity, absorbance signals were measured for both positive and negative controls of the RdRp reaction. Negative controls lacked the RdRp enzyme, while positive controls contained both the RNA substrate and the enzyme. The assay was conducted on four separate days to evaluate reproducibility. The results showed minimal variation in absorbance values across both control groups, with the majority of measurements falling within two standard deviations of the mean absorbance value and all the values falling within three standard deviations of the mean absorbance value (Figure 2).

**Figure 2.**
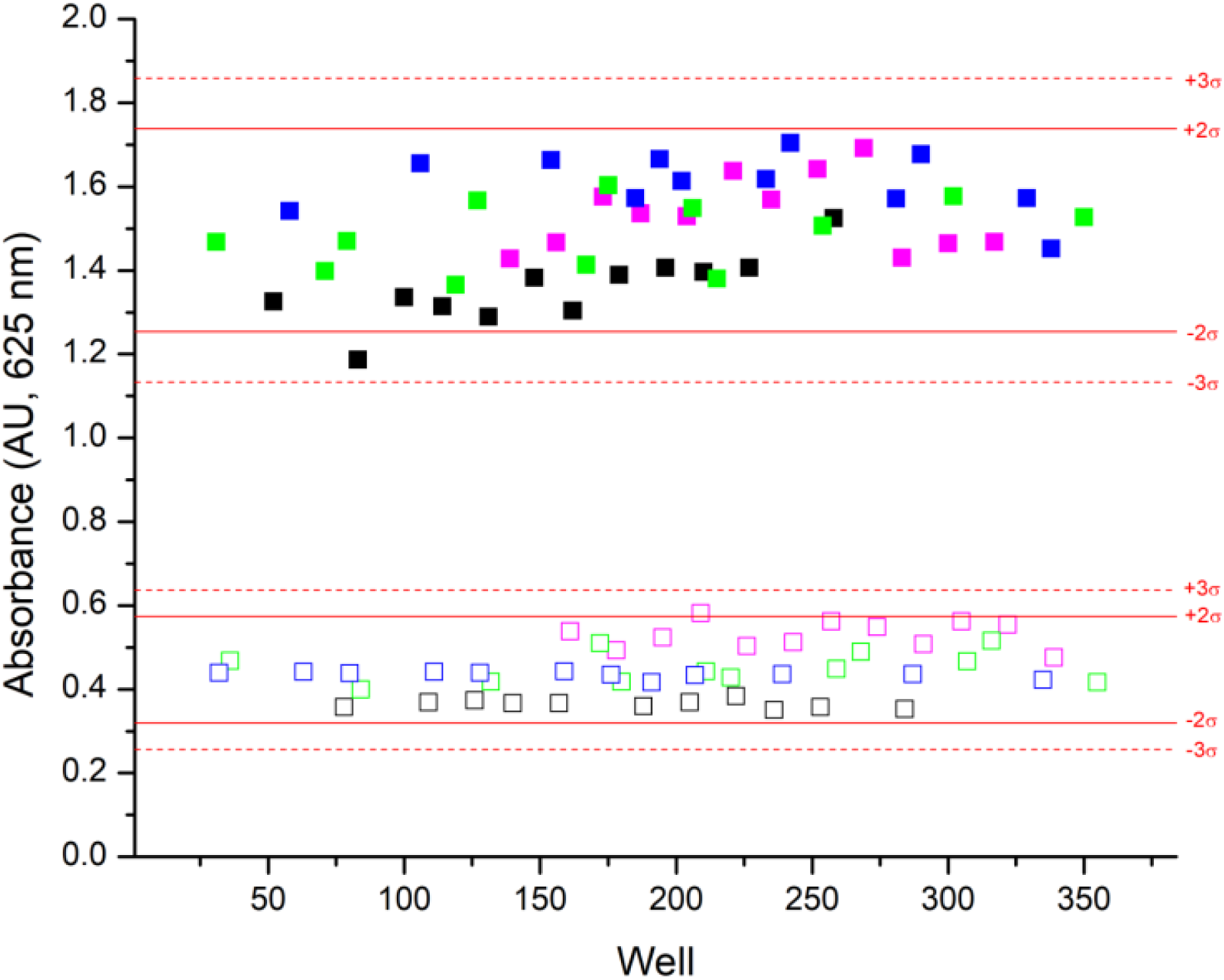
The malachite green–based assay employed in this study is reproducible across multiple days. Absorbance measurements were obtained for positive and negative controls using 384-well microplates across four independent assay days. Each color represents a different day (Days 1, 2, 3, and 4). Positive controls, containing ZIKV RdRp and RNA substrate, are represented by filled squares, and negative controls, lacking the enzyme, by empty squares. σ denotes the standard deviation from the mean absorbance values. The solid and dotted red lines indicate two and three standard deviations, respectively, from the mean of the positive or negative control groups.

The Z′ factor is a dimensionless statistical parameter used to assess the signal dynamic range and variability between the positive and negative controls in a HTS assay (Equation 1) ^17^. A high Z′ value indicates strong separation between positive and negative control signals and low data noise, both of which are essential for distinguishing true hits from experimental variability. An acceptable Z′ value ranges from 0.5 to 1 ^17^. All Z′ values determined from experiments conducted over four separate days fell within this range, indicating that the malachite green assay used here is robust, exhibits a high signal-to-noise ratio, and is suitable for HTS (Table 1).

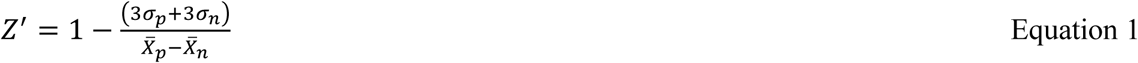

**Table 1.** The malachite green assay employed in this study is robust, has a large dynamic range, and a high signal-to-noise ratio. ^a^The malachite-green assay, with the negative and positive controls, was performed in different plates on four different days. The positive control contained both the RNA substrate and RdRp, while the negative control lacked the RdRp enzyme. ^b^Z′ values were calculated from experiments performed on four different days using Equation 1. The Z′ values indicate that the separation between the negative and positive controls is sufficient to reliably detect inhibitors. The Z′ values obtained across all four days of experimentation fall within the acceptable range of 0.5 < Z′ < 1, demonstrating that the assay exhibits high quality and is suitable for HTS ^17^. ^c^ SW was calculated using Equation 2 for four different plates across multiple days. The SW values are significantly larger than 2, the accepted SW value for an HTS-suitable assay ^17^.

| Plate/Day<sup>a</sup> | $Z'$<sup>b</sup> | SW<sup>c</sup> |
| --- | --- | --- |
| 1 | 0.731 | 9.115 |
| 2 | 0.655 | 7.796 |
| 3 | 0.666 | 8.790 |
| 4 | 0.807 | 13.911 |

In Equation 1, X̅p is the average malachite green assay absorbance of the positive control; X̅n is the average malachite green assay absorbance of the negative control; σp is the standard deviation of the positive control; and σn is the standard deviation of the negative control ^17^.

SW is a dimensionless parameter used to assess the difference in signal between negative and positive controls ^17^. SW was calculated using Equation 2 across four different days and plates (Table 1) ^17^. An SW value greater than 2 is considered adequate for detecting inhibitors in an HTS assay. The SW values obtained on different days were significantly greater than 2, demonstrating once again that our malachite green assay is both robust and sensitive ^17^.

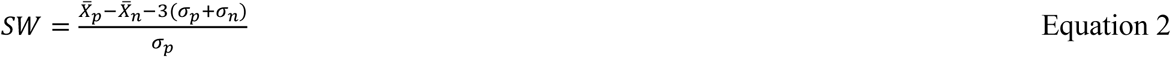

### Purpurogallin and Digallic Acid Inhibit NS5-RdRp Activity

The optimized green/molybdate colorimetric assay was used to screen for RdRp inhibitors. A total of 204 out of 419 natural products from the NCI Natural Products Set IV were screened. The screening was performed in the presence of 10μM of each compound (Table S1). We found that, among the 204 natural compounds investigated, two of the screened compounds—purpurogallin (National Service Center (NSC) 35676) and digallic acid (NSC 59263)—were effective in inhibiting the polymerase activity of ZIKV RdRp (Figure 3).

**Figure 3:**
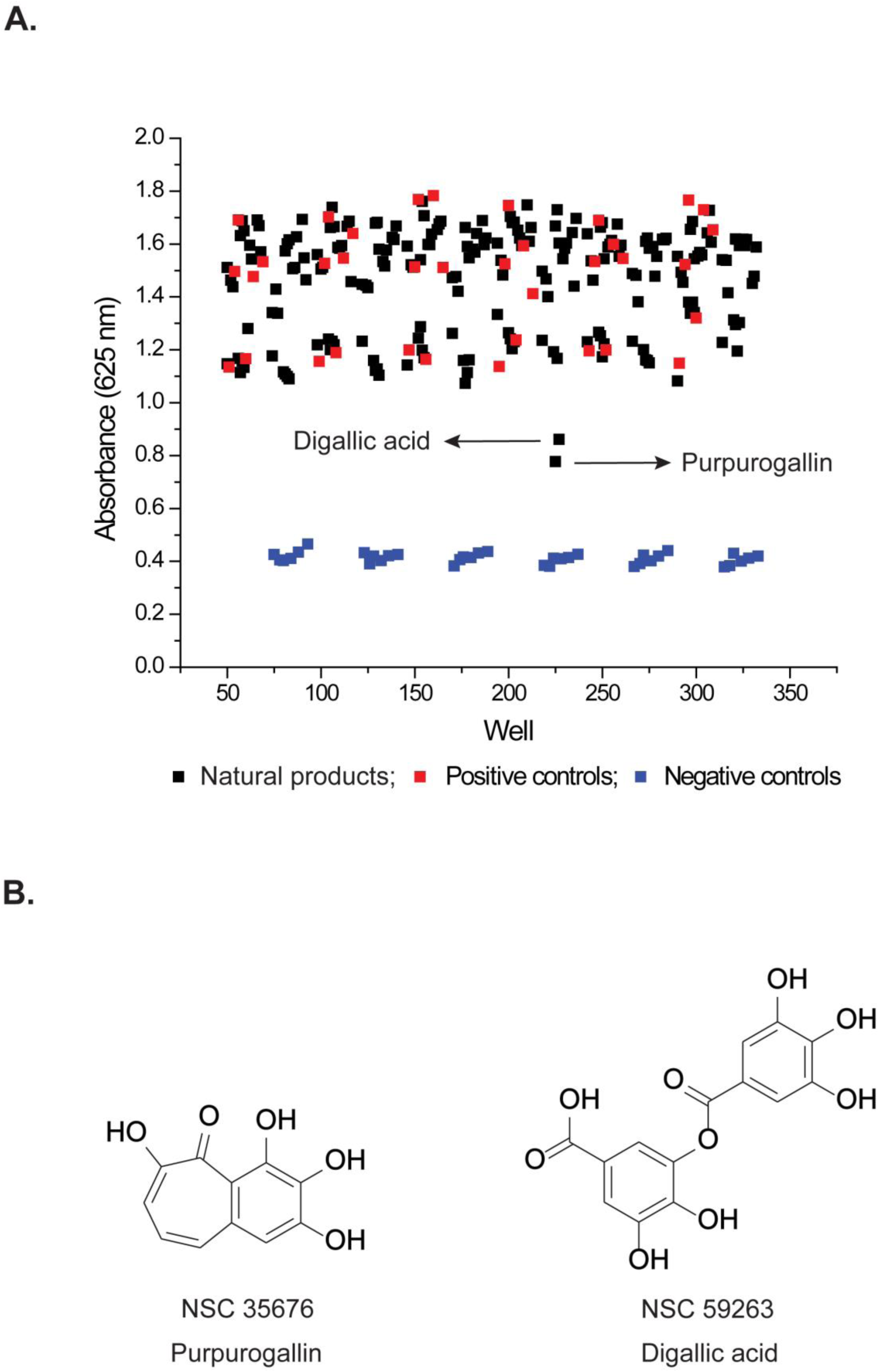
Purpurogallin and Digallic Acid Inhibit the Polymerase Activity of ZIKV RdRp as Determined by the malachite green assay. A) A total of 204 out of 419 compounds from the NCI Natural Products Set IV were screened using the malachite green assay. The list of natural products tested is provided in Table S1. The absorbance of the positive control reactions, which contained the RNA substrate and RdRp, is shown as red squares; that of the negative control reactions, which lacked the RdRp enzyme, as blue squares; and the absorbance of the reactions that contained one of the 204 compounds along with both RNA and RdRp as black squares. The two compounds—Purpurogallin (NSC 35676) and Digallic Acid (NSC 59263)—that inhibited the polymerase activity of RdRp are labeled. B) Chemical structures of the two compounds that inhibit the polymerase activity of RdRp: Purpurogallin (NSC 35676) and Digallic Acid (NSC 59263).

Purpurogallin, a benzotropolone derivative, occurs naturally in the nut galls of *Quercus* species or can be synthesized via pyrogallol oxidation ^24^. It has been shown to protect cells from hydrogen peroxide-induced DNA damage and apoptosis, both of which are associated with oxidative stress^24^. ZIKV infection induces oxidative stress, which facilitates viral replication ^25^. Therefore, purpurogallin may serve as a dual-function inhibitor by targeting ZIKV RdRp, an enzyme essential for viral replication, and by mitigating oxidative stress associated with infection.

Moreover, purpurogallin has demonstrated anti-inflammatory effects in human endothelial cells ^26–28^. Acute inflammation is considered a central mechanism underlying ZIKV-associated pathologies, and modulation of the cellular inflammatory response has been proposed as a promising therapeutic strategy ^2^. Importantly, the anti-inflammatory compound MK-591 has emerged as a potential candidate for the treatment of ZIKV infection ^29^. Accordingly, purpurogallin may function as a multifunctional inhibitor by attenuating the inflammatory response, inhibiting RdRp activity, and reducing oxidative stress.

Digallic acid, a compound derived from *Pistacia lentiscus* fruits or synthesized via partial hydrolysis of tannic acid, was shown in this study to inhibit ZIKV RdRp activity (Figure 3) ^30–31^. A previous study also reported that digallic acid inhibits the reverse transcriptase activity of murine leukemia virus (MLV) and human immunodeficiency virus (HIV) ^32^. In that study, digallic acid inhibited the activity of reverse transcriptase by competing for the template RNA binding site ^32^.

In addition to inhibiting MLV and HIV reverse transcriptase, digallic acid was shown to moderately inhibit the activity of human DNA polymerases alpha and beta, but did not inhibit human DNA polymerase gamma or *E. coli* DNA polymerase I ^32^. This cross-polymerase inhibition, combined with mechanistic studies, suggests that digallic acid may interact with conserved structural motifs involved in DNA or RNA template binding and shared among these human and viral enzymes. Future structural studies of ZIKV RdRp and HIV reverse transcriptase in the presence of digallic acid could aid in the design of inhibitors structurally similar to digallic acid, which are highly specific for ZIKV and HIV polymerases while sparing human DNA polymerases. These dual-acting inhibitors may hold therapeutic potential for managing co-infections caused by ZIKV and HIV.

## CONCLUSIONS

The optimized malachite green assay we employed here is robust, reproducible, and sensitive across different experimental days (Table 1). This assay, as optimized by us, could be employed for HTS of DNA and RNA polymerases, as well as other enzymes that produce pyrophosphate during their catalytic cycles. Through the screening of NCI natural product libraries, purpurogallin and digallic acid emerged as promising RNA-dependent RNA polymerase (RdRp) inhibitors (Figure 3, Table S1). Future experiments will validate these inhibitors using an orthogonal assay alongside the malachite green assay and determine their precise mode of action and potency both *in vitro* and in ZIKV-infected cell cultures.

## ASSOCIATED CONTENT

### Supporting Information

Complete list of NSC numbers for all compounds investigated in this study.

## ACKNOWLEDGMENTS

We are grateful to Luis Gracia Mazuca for the discussion regarding this manuscript.

## For Table of Content Use Only

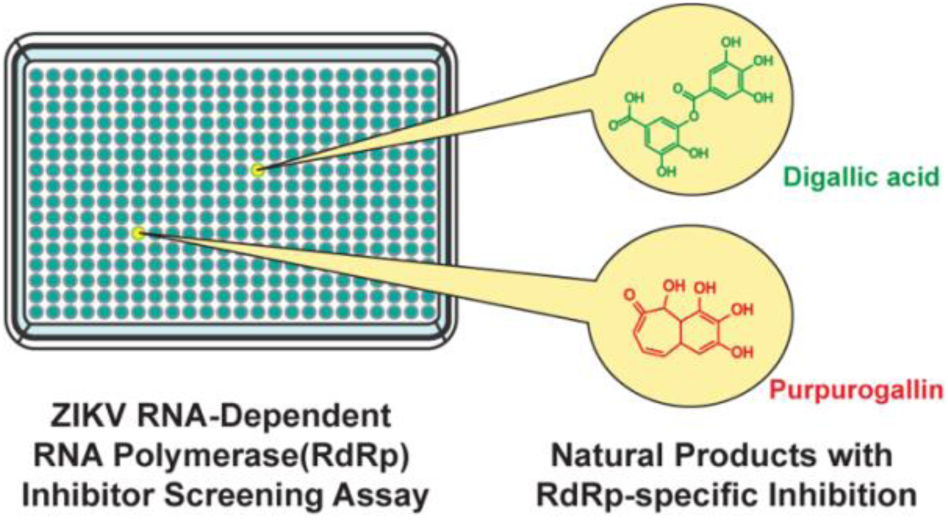

