## Supporting Information for "Discovery of Inhibitors Targeting the RNA-Dependent RNA Polymerase of the Zika Virus"

Table S.1. Complete list of National Service Center (NSC) numbers for all compounds investigated in this study.

| NSC[a] | | | | | |
|---|---|---|---|---|---|
| 757 | 56464 | 226080 | 637086 | 9699 | 31754 |
| 7524 | 71795 | 284200 | 2150 | 13123 | 34552 |
| 2952 | 58368 | 250429 | 661755 | 10105 | 32743 |
| 7668 | 72116 | 284437 | 2347 | 14135 | 34758 |
| 3053 | 60387 | 250430 | 719655 | 11866 | 32944 |
| 8519 | 76022 | 285116 | 2802 | 14664 | 35550 |
| 5113 | 62709 | 255109 | 824 | 11905 | 32984 |
| 9665 | 82151 | 287088 | 2835 | 14665 | 35676[b] |
| 5159 | 63701 | 263164 | 1115 | 12444 | 33410 |
| 11440 | 85236 | 292567 | 3071 | 14974 | 36294 |
| 5366 | 67574 | 270914 | 2080 | 12865 | 34202 |
| 11926 | 85239 | 301683 | 3590 | 15307 | 36354 |
| 12097 | 87511 | 302289 | 3716 | 15624 | 36407 |
| 23969 | 118343 | 332598 | 5897 | 19509 | 43338 |
| 13252 | 89671 | 305222 | 4143 | 16631 | 36437 |
| 26258 | 122023 | 332876 | 6435 | 19990 | 43339 |
| 14975 | 94600 | 307981 | 4586 | 18334 | 36508 |
| 26271 | 122224 | 333856 | 6832 | 20103 | 43871 |
| 15780 | 96911 | 325014 | 5036 | 18805 | 36693 |
| 26326 | 122750 | 337783 | 7521 | 20264 | 44138 |
| 22070 | 105388 | 325319 | 5379 | 19028 | 38010 |
| 29854 | 122819 | 339555 | 7525 | 21725 | 44175 |
| 22842 | 114344 | 330753 | 5863 | 19038 | 38270 |
| 30552 | 129536 | 345647 | 7532 | 21728 | 45384 |
| 31048 | 133100 | 349438 | 7533 | 22939 | 46709 |
| 36351 | 176503 | 382796 | 8661 | 26254 | 51351 |
| 31867 | 145118 | 350085 | 7535 | 23615 | 46728 |
| 36398 | 177406 | 400978 | 8751 | 26327 | 56410 |
| 32192 | 150817 | 361902 | 7606 | 23878 | 47147 |
| 42038 | 177858 | 401005 | 8797 | 27425 | 59258 |
| 32979 | 153858 | 369397 | 7616 | 24819 | 50131 |
| 45383 | 180515 | 407286 | 8973 | 28841 | 59263[b] |
| 32982 | 157035 | 375294 | 7652 | 24872 | 50132 |
| 45923 | 209870 | 407306 | 9170 | 30238 | 59729 |
| 35611 | 169627 | 376248 | 8625 | 24951 | 50393 |
| 51001 | 210236 | 614552 | 9248 | 30625 | 61809 |

[a] NSC is a unique identifier assigned by the NCI Developmental Therapeutics Program (DTP) to catalog and track chemical compounds. In this study, 204 compounds from the Natural Product Set IV of the NCI DTP were evaluated for their ability to inhibit Zika virus (ZIKV) RNA-dependent RNA polymerase (RdRp). This table lists the NSC numbers of all compounds investigated. More information about the compound plates is available at: https://wiki.nci.nih.gov/display/ncidtpdata/compound+sets. The complete chemical names of each compound can be found on the NCBI PubChem website: https://pubchem.ncbi.nlm.nih.gov/.
[b] The two compounds, purpurogallin and digallic acid, that inhibited ZIKV RdRp in the Malachite Green Assay are indicated in bold.
